# Specialized subsets of innate-like T cells and dendritic cells protect from lethal pneumococcal infection in the lung

**DOI:** 10.1101/2021.07.25.453697

**Authors:** Mallory Paynich Murray, Catherine M. Crosby, Paola Marcovecchio, Nadine Hartmann, Shilpi Chandra, Meng Zhao, Archana Khurana, Sonja P. Zahner, Björn E. Clausen, Fadie T. Coleman, Joseph P. Mizgerd, Zbigniew Mikulski, Mitchell Kronenberg

## Abstract

Innate-like T cells, including invariant natural killer T (iNKT) cells, mucosal-associated invariant T (MAIT) cells and γδ T cells, are present in various barrier tissues, including the lung. They carry out protective responses during infections, but the mechanisms for protection are not completely understood. Here, we investigated their roles during pulmonary infection with *Streptococcus pneumoniae*. Following infection, innate-like T cells rapidly increased in lung tissue, in part through recruitment, but TCR activation and cytokine production occurred mostly in IL-17-producing NKT17 and γδ T cells. NKT17 cells were preferentially located outside the vasculature prior to infection, as were CD103^+^ dendritic cells (cDC1), which were important both for antigen presentation to NKT17 cells and γδ T cell activation. Whereas IL-17A-producing γδ T cells also were numerous, GM-CSF was exclusive to NKT17 cells and contributed to iNKT cell-mediated protection. These studies demonstrate how particular cellular interactions and responses of functional subsets of innate-like T cells contribute to protection from pathogenic lung infection.

## Introduction

The lung is a barrier tissue and contains populations of innate lymphoid cells (ILC) and innate-like T cells. Innate-like T cells carry out rapid activation and cytokine secretion, thereby bridging the innate and adaptive immune responses. They have a number of distinct properties, including the ability to respond to TCR stimulation or to cytokines in the absence of TCR signaling (Gutierrez-Arcelus et al., 2019). These lymphocyte types include invariant natural killer (iNKT) cells, γδ T cells and mucosal-associated invariant T (MAIT) cells. iNKT cells respond to glycolipid antigens presented by CD1d, a non-polymorphic MHC class I antigen presenting molecule. A number of bacteria have been shown to produce glycolipids recognized by iNKT cells, including pathogens such as *Borrelia burgdorferi*, group B *Streptococcus*, and *Streptococcus pneumoniae* (Chang et al., 2011; Kinjo et al., 2011; Kinjo et al., 2006; Mattner et al., 2005). γδ T cells are not exclusively specific for microbial antigens, although the most abundant population of γδ T cells in human blood recognizes pyrophosphate containing microbial metabolites (Harly et al., 2012; Ribot et al., 2021; Sandstrom et al., 2014). MAIT cells recognize microbial riboflavin metabolites presented by MR1, a highly conserved and non-polymorphic MHC class I antigen presenting molecule. Many bacteria produce antigens that stimulate MAIT cells, including pathogens such as *Mycobacterium tuberculosis*, *Francisella tularensis* and *S. pneumoniae* (Hartmann et al., 2018; Le Bourhis et al., 2010; Meierovics et al., 2013).

Innate-like T cells differentiate into functional subsets in the thymus. For example, thymus iNKT cells differentiate into NKT1, NKT2 and NKT17 cells (Lee et al., 2013; Watarai et al., 2012), analogous in their capacity to produce cytokines to Th1, Th2 and Th17 cells, respectively. Functional subsets of γδ and MAIT cells that produce IFN-γ or IL-17 also are present in the thymus (Godfrey et al., 2015; Ribot et al., 2021). These populations likewise are found in peripheral tissues, including the lung, where they reside long-term (Khairallah et al., 2018; Murray et al., 2021; Salou et al., 2019; Thomas et al., 2011).

*S. pneumoniae* is a leading cause of pneumonia and meningitis in both children and the elderly (O’Brien et al., 2009; Wahl et al., 2018; Wroe et al., 2012). iNKT cell antigen-dependent responses to this microbe have been well characterized (Brigl et al., 2011; Girardi et al., 2011; Kinjo et al., 2011) and iNKT cells are essential for protection of mice from *S. pneumoniae* (Brigl et al., 2011; Kawakami et al., 2003; Kinjo et al., 2011). γδ T cells also have been shown to be important for protection from *S. pneumoniae*, with a role in the early recruitment of neutrophils (Hassane et al., 2017; Nakasone et al., 2007). Therefore, *S. pneumoniae* provides an example in which two types of innate-like T lymphocytes play non-redundant roles in protection from lung infection.

Although iNKT cells and γδ T cells are required for host defense, there is relatively little information on the mechanisms of activation, cell-cell contacts and immune responses underlying the role of innate-like T cells in host protection. Here, we addressed several issues pertaining to protective innate-like T cell responses to lung infection with *S. pneumoniae*. First, we sought to identify the underlying mechanisms leading to the rapid activation of protective iNKT and γδ T cell responses during *S. pneumoniae* infection, including the role of TCR activation of these cells. Second, we sought to identify a nonredundant function(s) of iNKT cells that would distinguish them from the more numerous γδ T cells that produce similar cytokines. Finally, we identified features of antigen presenting cell (APC) subsets important for activating iNKT cells and γδ T cells. Our data show how specialized responses of two populations of innate-like T cells, partially dependent on cDC1 activation and antigen presentation, provide host protection.

## Results

### Diverse *S. pneumoniae* strains have iNKT cell antigens

Previously, we reported that a URF918, a clinical isolate of an invasive serotype 3 strain of *S. pneumoniae*, has antigens that activate iNKT cells (Kinjo et al., 2011). To determine if different *S. pneumoniae* strains have antigens that activate iNKT cells, production of antigens was analyzed in an APC-free, hybridoma stimulation assay. In this assay, total bacterial sonicates were loaded onto CD1d coated plates and IL-2 secretion measured TCR-mediated activation. Antigenic activity was observed in the D39 strain (serotype 2), as well as in some strains from a group of clinical isolates of serotype 19A, including those that caused invasive infections (denoted by *, Supplementary Figure 1, Supplementary Table 1). Therefore, these data suggest that diverse serotypes of *S. pneumoniae* are likely to have antigens that activate iNKT cells. There was variability in antigenic content, however, as we observed previously for MAIT cell antigens in these strains (Hartmann et al., 2018). We used URF918 for the following experiments, because as shown previously, when C57BL/6J mice were infected with this strain bacterial clearance and survival were highly dependent on the activation of iNKT cells (Brigl et al., 2011; Kawakami et al., 2003; Kinjo et al., 2011).

### Vasculature and tissue location of innate like lung T cells

We analyzed the extent to which individual iNKT cell subsets, γδ T cells and MAIT cells were located in the vasculature, before and early after infection with *S. pneumoniae*. iNKT cell subsets were enumerated by flow cytometry using *α*GalCer-loaded CD1d tetramers together with iNKT cell subset-specific cell surface proteins (Supplementary Figure 2) that have been shown to provide highly enriched populations (Engel et al., 2016; Murray et al., 2021; Zhao et al., 2018). At steady-state, the three prevalent iNKT cell subsets in the lungs were NKT1, NKT17, and NKT2 cells, in order of decreasing prevalence (Figure 1A). The number of lung iNKT cells significantly increased by 15 hours and further at 24 hours after infection, with the largest increase in NKT1 cells (Figure 1A). This was primarily due to recruitment, rather than cell proliferation, as shown by comparable BrdU incorporation in iNKT cells from mice analyzed at 24 hours after infection (Figure 1B). Despite increased BrdU incorporation, NKT2 cells remained a relatively minor subset after infection.

**Figure 1:**
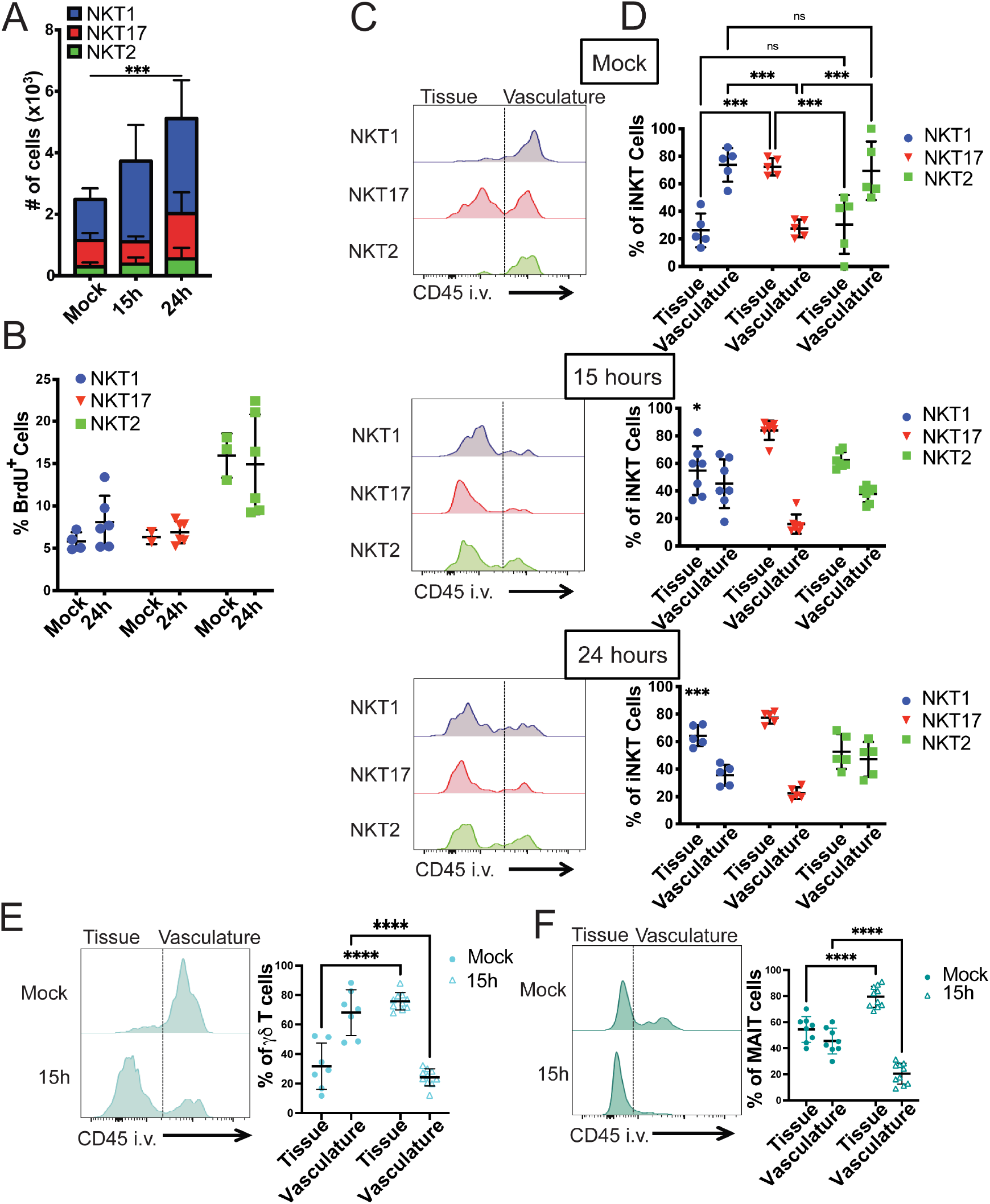
Innate-like T cells are recruited into the lung tissue after *S. pneumoniae* infection. A) iNKT cell subset numbers in the lungs before and after infection. N=7 mice per group, 2 independent experiments combined. Statistical significance was assessed via 2-way ANOVA, with Tukey’s multiple comparisons test. Source of variation: Interaction (*); Time (***), depicted in figure; Subset (****). NKT1 cells significantly expand at 15h (**) and 24h (****), Tukey’s multiple comparisons test. B) Percent of BrdU^+^ iNKT cell subsets in lung 24h after infection. N=6 mice per group, combined data from 2 independent experiments. C) Representative histograms of sub-tissue location of lung iNKT cell subsets, as determined by anti-CD45 antibody intravenous injection prior to tissue collection. Line separating tissue and vasculature based on iNKT cell subsets from mice not injected with anti-CD45. D) Quantification of (C). N=7 mice per group, combined data from 3 independent experiments. Statistical significance of NKT subset localization in Mock assessed via 2-way ANOVA, with Šídák’s multiple comparisons test. Tissue NKT1 mock vs. tissue NKT1 15h, p = 0.0113 (*); Tissue NKT1 mock vs. tissue NKT1 24h, p = 0.0004 (***), unpaired t test. E &F) Representative histograms and quantitation of sub-tissue location of lung γδ T cells and MAIT cells, as determined by anti-CD45 antibody intravenous injection prior to tissue collection. Combination of 2 independent experiments, N = 7-10 mice per group. Statistical significance assessed via 2-way ANOVA, with Šídák’s multiple comparisons test.

We analyzed the location of the iNKT cell subsets by harvesting tissues 3 minutes after labeling with an anti-CD45 antibody. This labels CD45^+^ cells located in the vasculature, and leaves cells in the interstitial tissue and alveoli unlabeled (Barletta et al., 2012; Reutershan et al., 2005). In agreement with a previous report (Salou et al., 2019), NKT17 cells had a significant presence in the lung, an average of more than 70% in the tissue, and this tendency was maintained or increased after infection (Figure 1C, D). In contrast, prior to infection, the majority of NKT1 and NKT2 cells were in the lung vasculature. Within 15 hours of infection, however, a higher proportion of NKT1 and NKT2 cells were no longer in blood vessels (Figure 1C, D). Similarly, the majority of the lung γδ T cells were also in blood but were recruited to the tissue by 15 hours (Figure 1E). Although MAIT cells were predominately located within the tissue at steady-state, the percentage of MAIT cells outside the vasculature also increased following infection (Figure 1F).

### Differential activation of subsets

We determined if subsets of lung iNKT cells were activated and produced their signature cytokines. Previous research showed that iNKT cells produce cytokines as early as 12 hours after infection (Holzapfel et al., 2014; Kinjo et al., 2011). To measure T cell antigen receptor (TCR)-mediated activation, we immunized Nur77^GFP^ mice, which express GFP under the control of the *Nr4a1* (Nur77) promoter (Moran et al., 2011). Reporter expression has been shown to be faithful readout of TCR activation in T lymphocytes, including iNKT cells (Holzapfel et al., 2014; Moran et al., 2011). We examined GFP expression in reporter mice by immunofluorescence staining of whole-mount vibratome sections of lung tissue from mice infected 14h earlier (Figure 2A). Although iNKT cells were infrequent, and therefore not present in every field, some cells stained brightly with the CD1d tetramer. Only duller staining was observed in sections stained with an unloaded CD1d tetramer. Many of the CD1d-tetramer^+^ cells also were positive for the GFP reporter, suggesting that some iNKT cells received a TCR signal following infection. Blood and lymphatic vessels express CD31 and iNKT cell staining in relation to CD31 staining suggests that most iNKT cells were outside of vessels following infection.

**Figure 2:**
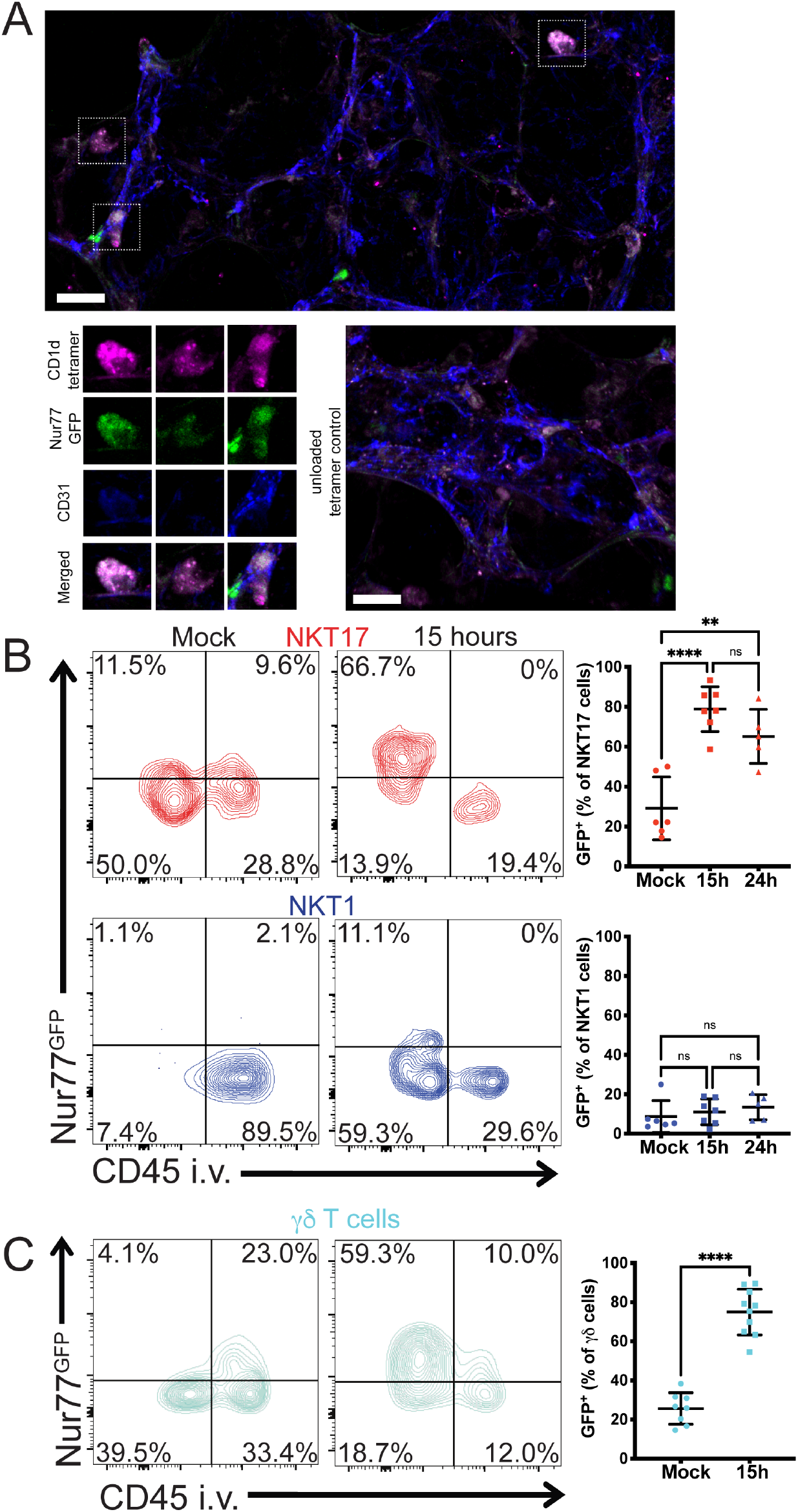
NKT17 and γδ T cells receive TCR signals after infection. A) Immunofluorescence staining of lung tissue in Nur77^GFP^ reporter mice 14 hours post-infection. CD31 (blue): blood vessels; CD1d-*α*GalCer-loaded tetramer (purple): iNKT cells; GFP (Green) indicating TCR reporter expression. Top: boxes indicate iNKT cells. Lower left: magnified view of the cells above, including two that are GFP^+^ and one that is not. Lower right: unloaded CD1d tetramer staining. Scale bar: 20 µm. B) Left: representative flow cytometry plot of sub-tissue location of TCR-activated (*Nur77*^GFP+)^ lung NKT17 and NKT1 cells at 15 hours after infection. Right: Quantification of GFP^+^ cells of panel A at 15 and 24 hours after infection. N= 5-7 mice per group, 3 independent experiments. **p=.0014, ****p<0.0001 (One-way ANOVA with Tukey’s multiple comparisons). C) Left: representative flow cytometry plot of sub-tissue location of TCR-activated (*Nur77*^GFP+)^ lung γδ T cells at 15 hours after infection. Right: Quantification of GFP^+^ cells at 15 hours after infection. N= 8-10 mice per group, combined data from 3 independent experiments. ****p<0.0001 (unpaired t test).

To achieve quantitative data, we used flow cytometry and focused on the more abundant lung subsets, NKT1 and NKT17 cells. There was a significant GFP signal in unstimulated NKT17 cells, consistent with previous data indicating that a minority of lung iNKT cells showed signs of constitutive activation (Murray et al., 2021). By 15 hours, the majority of NKT17 cells were GFP^+^, and these reporter positive cells were inaccessible to the CD45 antibody and therefore were outside the vasculature (Figure 2B). Very few NKT1 cells were GFP^+^ and the mean fluorescence intensity of the few GFP^+^ cells was lower, although these TCR-activated NKT1 cells were also unlabeled by anti-CD45 (Figure 2B). Therefore, NKT17 cells were the predominant iNKT cell subset receiving a TCR signal at early times after infection, and TCR responses correlated with an extravascular location. An increased percentage of γδ T cells also became GFP^+^ after infection, although like NKT17 cells, they also showed evidence for constitutive activation (Figure 2C).

### Role of cytokines in protection

We assessed cytokine production by innate-like T cells following *S. pneumoniae* infection by intracellular cytokine staining. Cells were analyzed ex vivo after a brief culture, without further TCR re-stimulation. At 15 hours post-infection, a substantial fraction of total iNKT cells produced IL-17A while relatively few produced IFN-γ (Figure 3A). In some experiments, we found IFN-γ^+^ NKT1 cells, but the percentage of expressing cells was never comparable to IL-17^+^ NKT17 cells (Figure 3B, Supplementary Figure 3A). The NKT17 cell population expressed both IL-17A and GM-CSF (Figure 3B), and included double (IL-17^+^, GM-CSF^+^) and single producers of one of these cytokines. NKT1 cells did not produce GM-CSF or IL-17A (Figure 3B). As previously reported, we confirmed that both IL-17A and GM-CSF were required for protection from *S. pneumoniae* infection (Brown et al., 2017; Ivanov et al., 2012; Zhang et al., 2009), with mice deficient in either of these cytokines displaying an increase in bacterial burden in the lung (Supplementary Figure 3B).

**Figure 3.**
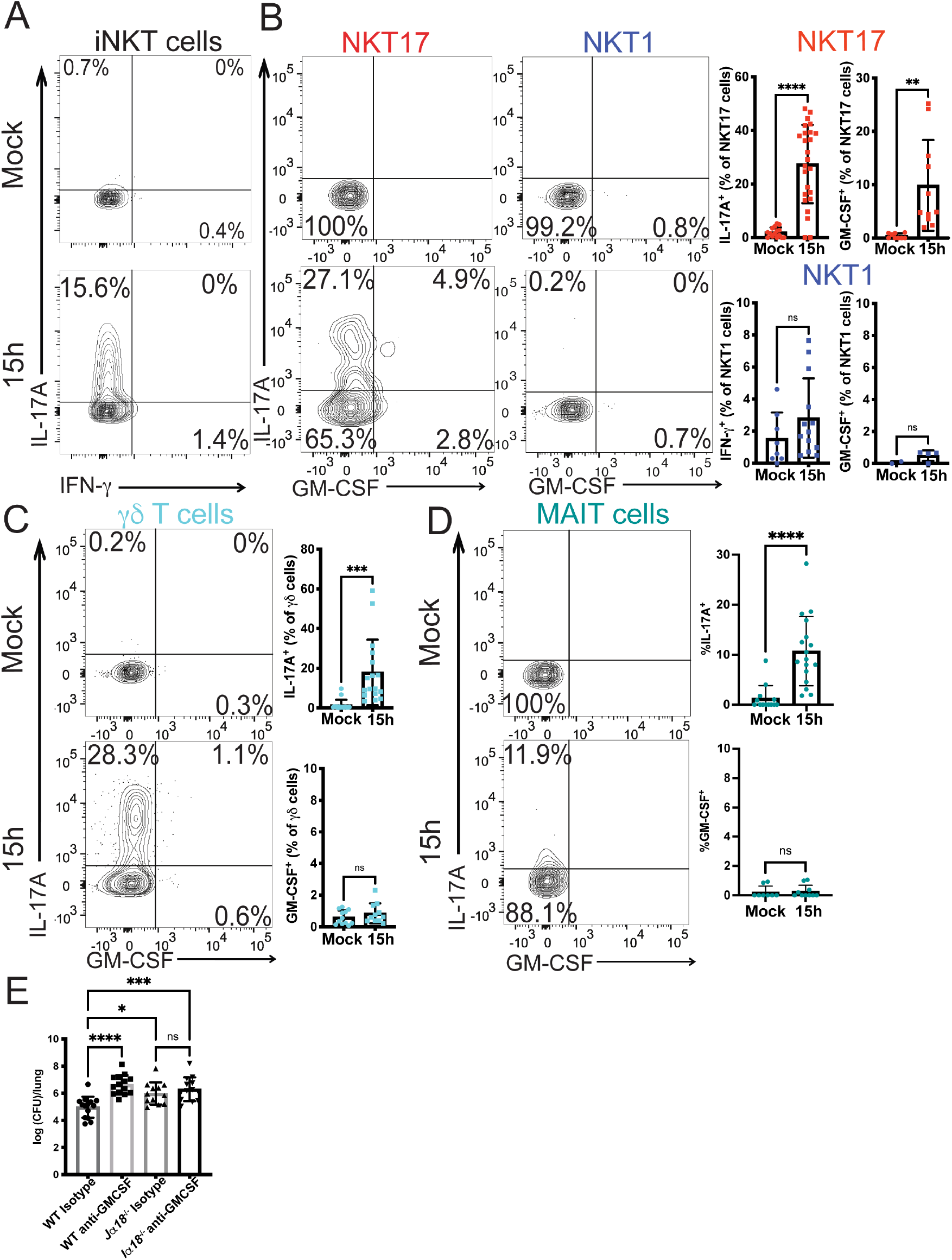
Innate-like T cells produce cytokines following infection. A) Representative plot of cytokine production by lung iNKT cells at 15 hours after infection with URF918. B) Left: Representative plot of cytokine production by NKT17 and NKT1 cells 15 hours after infection with URF918. Right: quantification of cytokine production by lung NKT1 cells and NKT17 cells. IL-17A, IFN-γ: N=8-25 mice per group, 5 independent experiments. GM-CSF: N=9-11 mice per group, representative data of 4 independent experiments; *p=.0038, ****p<0.0001 (unpaired t test). C & D) Cytokine production by γδ T cells (C) and MAIT cells (D) 15 hours post infection with URF918. Representative flow cytometry plots on left, quantitation on right. C) N=11-17 mice per group, combined from 4 independent experiments, *** p = 0.0004, unpaired t test. D) IL-17A: N=13-17 mice per group, combined from 4 independent experiments; GM-CSF: N=6-10 mice per group, 2 independent experiments, **** p<0.0001, unpaired t test. E) Bacterial burdens in lung (16 hpi) of C57BL/6J mice and *Jα18*^-/-^ mice treated with an isotype control (Rat IgG2a) or anti-GM-CSF antibody and infected with URF918.

γδ T cells and MAIT cells also rapidly increased IL-17A synthesis following infection, but neither cell type produced GM-CSF in response to *S. pneumoniae* infection (Figure 3C, 3D) (Hassane et al., 2020; Ivanov et al., 2014). MAIT cells were relatively infrequent, and analysis of *Mr1^-/-^* mice indicated they were not essential for protection (Supplementary Figure 3C). IL-17A-producing γδ T cells were more numerous after *S. pneumoniae* infection than iNKT cells (approximately 15,000 γδ T cells compared to less than 1,000 NKT17 cells following infection), suggesting that IL-17A production alone by iNKT cells would not account for their requirement for host protection.

Considering the strong effect on susceptibility of GM-CSF deficiency (Supplementary Figure 3B), the importance of neutrophil recruitment for defense from *S. pneumoniae* (Garvy and Harmsen, 1996), and the early production of GM-CSF uniquely by iNKT cells, these data suggest that production of GM-CSF may be one factor responsible for the non-redundant function of iNKT cells. To determine if iNKT cell GM-CSF was required, we treated C57BL/6J mice and *Jα18*-deficient mice, lacking iNKT cells, with an anti-GM-CSF antibody prior to infection. If GM-CSF was required for iNKT cell-mediated protection, then blockade of GM-CSF in the absence of iNKT cells should have no effect on bacterial burden. Indeed, we found that C57BL/6J mice treated with an anti-GM-CSF antibody had an increased bacterial burden following infection compared to isotype-treated C57BL/6J control mice (Figure 3E). *Jα18*-deficient mice treated with the anti-GM-CSF or isotype control antibody, however, had similar bacterial burdens (Figure 3E), suggesting that iNKT cell production of GM-CSF contributed to protection from infection.

### Differential responses of lung DC subtypes to infection

The lung contains several types of APCs that could participate in iNKT cell activation during infection. As shown in Figure 4A, prior to infection, cDC2 cells, gated as in Supplementary Figure 4A, were in both the lung tissue and vasculature, while the less numerous CD103^+^ or cDC1 cells were almost exclusively outside the vasculature and presumably within the tissue. 15 hours after infection, there was a significant increase in cDC2 cells in both the tissue and vasculature. There also was an influx of CD64^+^ monocyte-derived dendritic cells (moDCs) into the lung tissue, which were rare in uninfected mice. All three DC subsets and alveolar macrophages expressed surface CD1d and thus could have the capacity to present antigen to iNKT cells (Supplementary Figure 4B).

**Figure 4:**
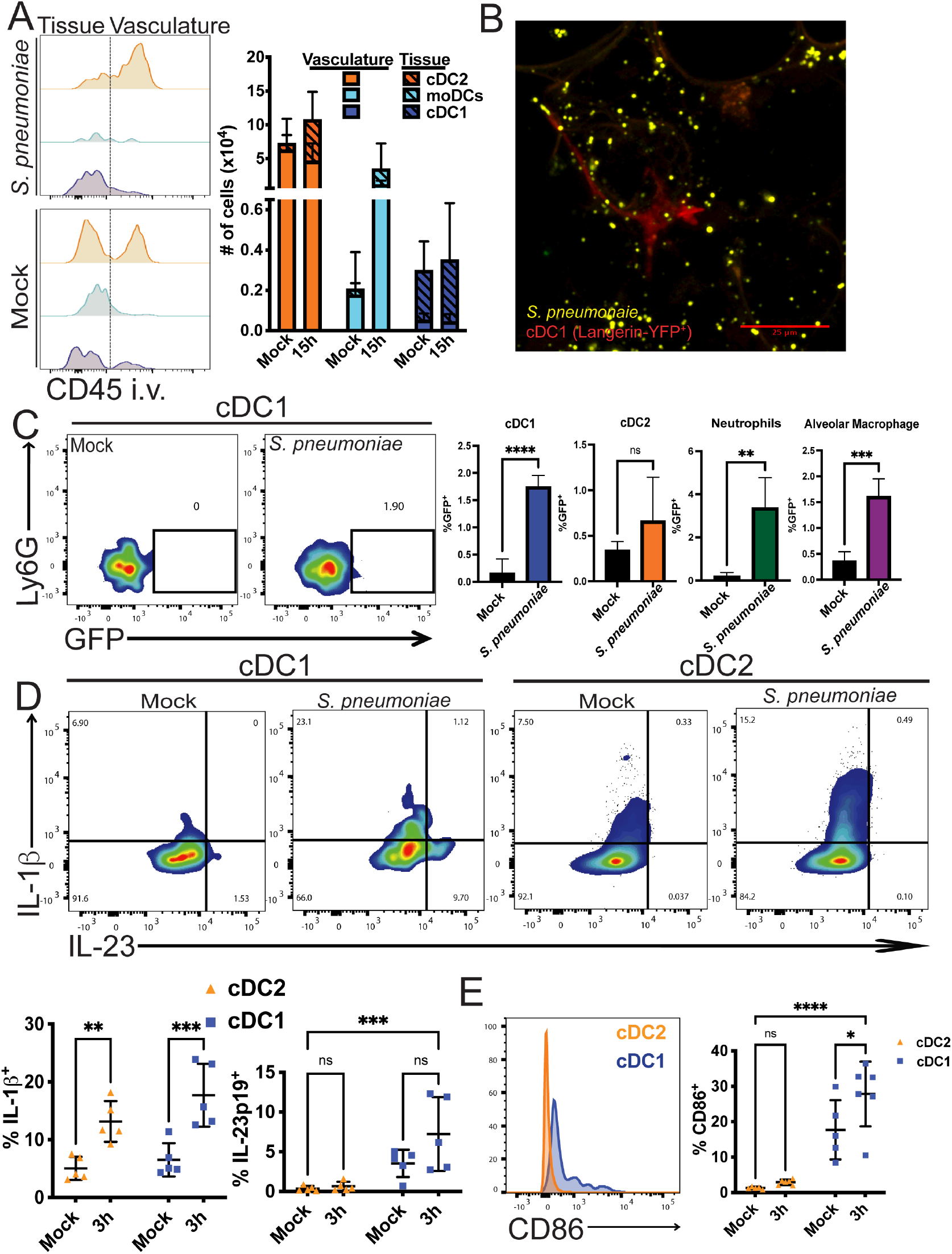
DC responses in the lung following infection. A) Left: representative histograms of sub-tissue location of dendritic cell subtypes at 0 and 15 hours after infection. Line depicting tissue vs. vasculature localization drawn based on each cell type from mice not injected with anti-CD45. Right: Quantification of cells outside of vasculature (hashed bars) and lung vasculature (open bars) at 0 and 15 hours after infection. N = 11-15 mice per group, 4 independent experiments combined. B&C) Mice were infected with URF918 and euthanized 2 hpi. B) GFP-expressing URF918 bacterial uptake in Langerin-YFP^+^ mice by immunofluorescence. C) Bacterial uptake by lung APCs was assessed via flow cytometry. Left: Representative flow cytometry of cDC1. Right: Quantification. N = 3-6 mice per group, combined from 3 independent experiments; statistical significance assessed via unpaired t test. D) Representative flow cytometry plot of cytokine production by lung cDC1 and cDC2 cells 3 hpi with URF918. Quantification below. Statistical significance was assessed via 2-way ANOVA, with Šídák’s multiple comparisons test. E. CD86 expression by cDC1 and cDC2 cells 3 hpi with URF918. Quantification on right. D & E. N = 5 mice per group, 2 independent experiments. Statistical significance was assessed via 2-way ANOVA, with Šídák’s multiple comparisons test.

To identify APC that could be important for defense from *S. pneumoniae*, we tested the capacity of lung myeloid cell types to take up bacteria. Mice were infected with GFP-expressing *S. pneumoniae* and 2 hours post-infection, bacterial uptake was assessed by immunofluorescence and flow cytometry. For immunofluorescence analysis, we analyzed Langerin-YFP mice in which cDC1 cells express YFP (Ghigo et al., 2013; Zahner et al., 2011). In tissue sections of infected Langerin-YFP mice, some *S. pneumoniae* puncta (yellow pseudo color) were co-localized with cDC1 cells (red pseudo color) (Figure 4B). By flow cytometry, we found a significant increase in GFP^+^ cDC1 cells (Figure 4C), as well as GFP^+^ neutrophils and alveolar macrophages. These data suggest that several myeloid cell types rapidly take up bacteria following infection.

iNKT cell production of IL-17 may be enhanced by either IL-23 or IL-1β, alone or in combination (Doisne et al., 2011; Rachitskaya et al., 2008; St Leger et al., 2018). Further, IL-23 has been shown to stimulate GM-CSF production by CD4^+^ Th17 cells, although it is not known if this is true for iNKT cells (Codarri et al., 2011; El-Behi et al., 2011; Rothchild et al., 2014). We therefore analyzed cytokines and cell surface molecules expressed by cDC1 and cDC2 that might be important for amplifying iNKT cell protective responses (Figure 4D). Pro-IL-1β production was increased in both cDC2 and cDC1 cells at 3 hours after infection (Figure 4D). IL-23 has been shown to be important for defense against *S. pneumoniae* (Kim et al., 2013) and IL-23p19 production was produced mainly by cDC1 cells. These cells also expressed much higher levels of the costimulatory molecule CD86 (Figure 4E). Due to their localization outside the vasculature, CD1d expression, bacterial uptake, cytokine production and co-stimulatory molecule expression, these data suggest that cDC1 may be an important subset required to stimulate the protective anti-microbial response in the lung.

### cDC1 activate lung T cells

The bacterial load was reduced in *Cd1d1^f/f^* x *CD11c*-Cre^+^ mice that lack CD1d in all dendritic cells (Supplementary Figure 4C). To address the hypothesis that cDC1 are most important, we infected *Batf3^-/-^* mice, which lack these cells due to deletion of an essential transcription factor (Grajales-Reyes et al., 2015; Hildner et al., 2008; Seillet et al., 2013). When *Batf3^-/-^* mice were infected with *S. pneumoniae*, iNKT cells did not expand and there was a 50% reduction in the frequency of IL-17A producing NKT17 cells compared to C57BL/6J mice (Figure 5A, B), similar to *Cd1d1^f/f^* x *CD11c*-Cre^+^ mice (Supplementary Figure 4C). This directly correlated with a reduction in neutrophil recruitment at 15 hours (Figure 5C), and an increase in bacterial burdens after 2 days (Figure 5D), altogether suggesting cDC1 cells are important for iNKT cell activation and protection from *S. pneumoniae*. Analysis of *Batf3^-/-^* mice revealed that expansion of γδ T cells, as well as cytokine production by these cells, was also dependent on cDC1 cells (Figure 5E).

**Figure 5:**
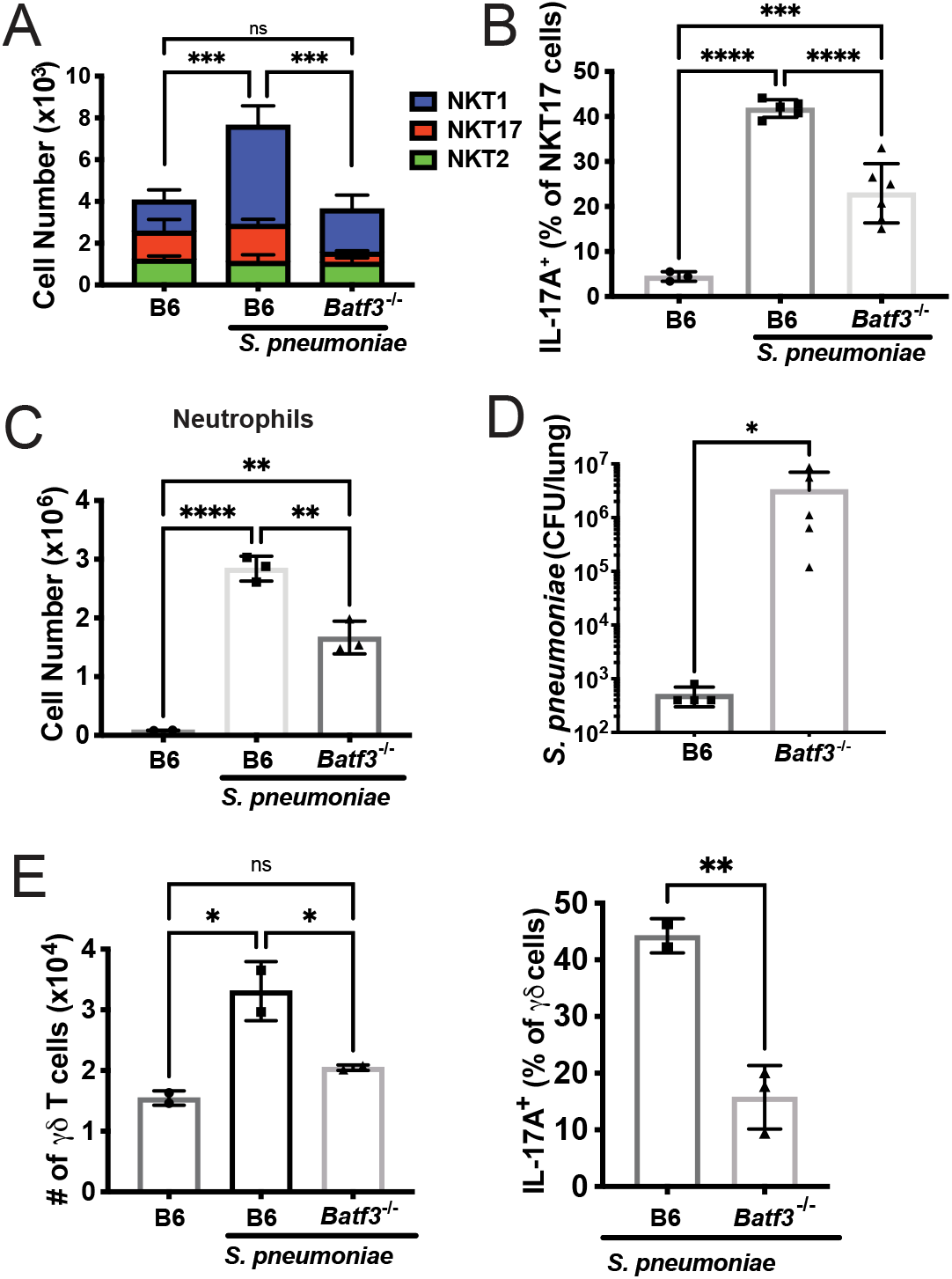
*Batf3^-/-^* mice have reduced innate-like T cell responses. A) iNKT cell numbers at 15 hours after infection in C57BL/6J and *Batf3*^-/-^ mice. N=6 per group. ***p<0.001, One-way ANOVA, with Tukey’s multiple comparisons. B) IL-17A production by NKT17 cells at 15 hours after infection. N=6 per group. ***p=.0004, ****p<0.0001 (One-way ANOVA with Tukey’s multiple comparisons) C) Lung neutrophil numbers at 15 hours after infection. N=6 per group. ****p<0.0001. D) Bacterial loads at 2 days. N=5 per group. *p=0.0159 (Mann-Whitney Test). E) γδ T cells numbers (left) and IL-17A production (right) at 15 hours after infection in C57BL/6J and *Batf3*^-/-^ mice. N=2-3 mice per group, 1 experiment. Left: One-way ANOVA with Tukey’s multiple comparisons; Right: ** p= 0.0078, unpaired t test.

To determine if protection was dependent on direct stimulation of iNKT cells and/or γδ T cells by antigens presented by CD1d, we utilized mice with conditional deletion of *Cd1d1* on cDC1 cells (*Langerin*-Cre^+^) (Zahner et al., 2011). The specificity of the deletion on populations of lung myeloid cells is shown in Supplementary Figure 4B. Deletion of *Cd1d1* had no effect on iNKT cell recruitment (Figure 6A, left), suggesting that innate immune activation of cDC1 and other cell types likely leads to the synthesis of chemokines that recruit iNKT cells in the absence of antigen presentation. Loss of CD1d expression by cDC1 cells significantly reduced iNKT cell-derived IL-17A (Figure 6A, center), however, and led to increased bacterial loads at 2 days (Figure 6A, right). Therefore, cDC1 cells not only participated in recruiting iNKT cells in a CD1d-independent fashion, but they activated these lymphocytes through CD1d antigen presentation. In contrast, IL-17A production by γδ T cells was not reduced in mice lacking CD1d specifically on cDC1 cells (Figure 6B), and therefore TCR activation of these cells must have depended on different antigens that were not dependent on CD1d.

**Figure 6:**
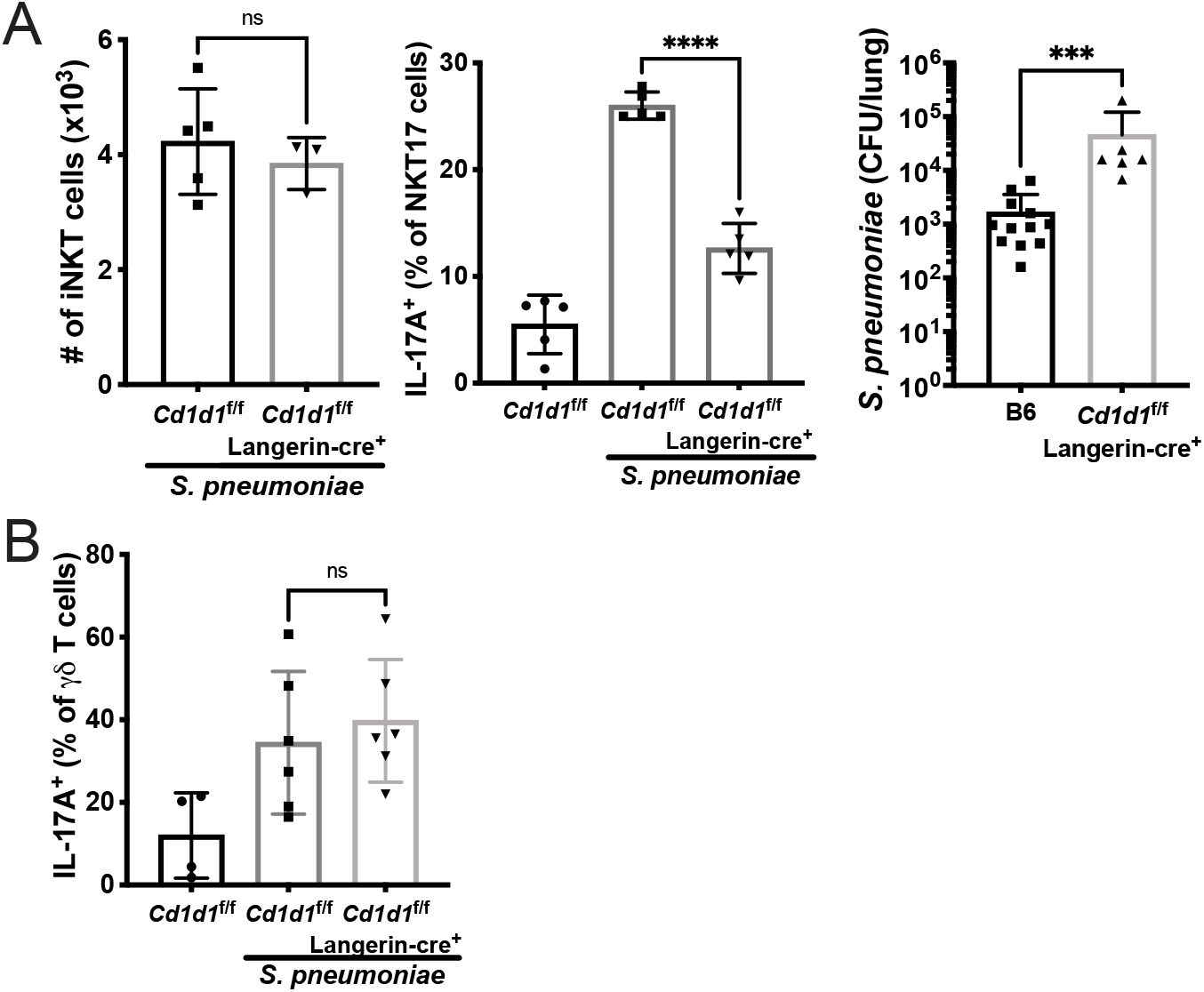
Dendritic cells deficient in CD1d have reduced iNKT cell responses. A) Langerin^cre^*Cd1d*^fl/fl^ mice following infection with URF918. Left: total iNKT cell numbers at 15 hours post-infection. N=5 per group. Middle: IL-17A production by NKT17 cells at 15 hours after infection. N=5 per group. ****p<0.0001 (One-way ANOVA with Tukey’s Multiple Comparisons). Right: Bacterial loads 2 days after infection. ***p<0.0001 (Mann-Whitney Test). B) IL-17A production by γδ T cells in *Cd1d1^f/f^* x *Langerin*-Cre^+^ mice following URF918 infection.

## Discussion

Here we have identified the cellular interactions and the means by which innate-like T cells participate in protection from lung infection with an important pathogen, *S. pneumoniae*. The different serotypes of *S. pneumoniae* that have been tested all have an antigen(s) that activates iNKT cells, but the requirement in vivo for a response by iNKT cells for host defense may be restricted to those strains that are more virulent (Barthelemy et al., 2016; Ivanov et al., 2012; Kawakami et al., 2003; Kinjo et al., 2011). Therefore, lung infection of mice with strain URF918 *S. pneumoniae* has provided a striking example in which the response of iNKT cells has proven to be absolutely essential for controlling CFU, beginning as early as 24h (Brigl et al., 2011; Kawakami et al., 2003; Kinjo et al., 2011). In this case, the protective role of iNKT cells is elicited very rapidly, with kinetics similar to features of the innate immune response. In contrast, although *S. pneumoniae* has an antigen that activates MAIT cells (Hartmann et al., 2018), and they produced cytokines rapidly after infection, we did not find an essential role for MAIT cells, perhaps due to their lower cell numbers in the lung. γδ T cells also are essential for host defense (Hassane et al., 2017; Nakasone et al., 2007), however, and they also are activated within 16h to produce cytokines such as IL-17 (Hassane et al., 2020). Previous research showed IL-17 could be detrimental or protective, depending on the infecting *S. pneumoniae* strain (Ritchie et al., 2018). Regardless, the early production of IL-17 by γδ T cells is correlated with the recruitment of neutrophils, a critical feature of the protective response (Hassane et al., 2017; Hassane et al., 2020). The requirement for both innate-like T cell types raises the issue as to how the contributions of iNKT cells and γδ T cells differ.

Consistent with a requirement for early responses, lung iNKT and γδ T cells are present in the lung and in a partially activated state (Murray et al., 2021). Furthermore, the number of both cell types rapidly increased in the lung after infection. Although the majority of both iNKT and γδ T cells were in blood vessels prior to infection, they were found predominantly outside the vasculature, and likely in the tissue parenchyma, by 15h. For iNKT cells, however, there is evidence for a TCR signal even earlier, by 5h (Holzapfel et al., 2014), and intracellular cytokine could be detected directly ex vivo by 12h (Holzapfel et al., 2014; Kinjo et al., 2011). Therefore, despite their rapid recruitment from the lung vasculature, some of the very earliest and perhaps critical responses by iNKT cells may be carried out by those iNKT cells already residing outside the vasculature. Lung vascular localization prior to infection, however, was not an impediment for rapid TCR activation of lung γδ T cells. Furthermore, there was evidence for constitutive activation of some γδ T cells, perhaps by products derived from the commensal microbiota.

Following *S. pneumoniae* infection, γδ T cells produce IFNγ and IL-17A (Hassane et al., 2020), and as noted, their role in protection has been linked to IL-17A production (Hassane et al., 2017; Hassane et al., 2020). Like γδ T cells, iNKT cells have been shown to participate in the protective response to a variety of infections, but in the majority of cases, protection was related to IFN-γ secretion by these cells (Kinjo et al., 2013). Even in other contexts, such as amelioration of arthritis, NKT1 cells were shown to be the effective population, which may in part reflect the prevalence of these cells in C57BL/6J mice (Zhao et al., 2018). Here our evidence suggests that the NKT17 cells are uniquely important for protection. The population of iNKT cells prior to infection outside the lung vasculature were predominantly NKT17 cells, in agreement with earlier reports (Lee et al., 2015; Salou et al., 2019). Therefore, tissue residence and early immune responses by NKT17 cells were correlated. Lung NKT1 cells are not exhausted, however, because after priming with *α*GalCer, they produced IFN-γ which protected from infection with *S. pneumoniae* along with IL-17A (Ivanov et al., 2012). Furthermore, in several earlier studies (Brigl et al., 2011; Holzapfel et al., 2014; Ivanov et al., 2012; Kinjo et al., 2011; Nakamatsu et al., 2007) there was evidence for IFN-γ production by iNKT cells after *S. pneumoniae* infection, although the percentage of cytokine-producing cells varied. We do not discount the importance of NKT1 cells and IFN-γ in our mice. The influx of NKT1 cells recruited to the lung and NKT1 responses later than 24h could have protective roles.

Considering that the capacity for IL-17A synthesis is found in several types of innate or innate-like cells, including ILC3 (Cua and Tato, 2010; Rosine and Miceli-Richard, 2020) as well as T cells, there could be functional redundancy for these cell types. In agreement with a previous publication (Hassane et al., 2020), we found that lung γδ T cells were more numerous than iNKT cells, and a higher percentage of them produced IL-17A compared to iNKT cells after *S. pneumoniae* infection. Therefore, it seemed unlikely that the nonredundant function of iNKT cells was due solely to IL-17A secretion, although differences in neighboring cells, combined with differences in other signals, could provide for a divergence in the effects of iNKT and γδ T cell IL-17. Some NKT17 cells in fact produce other cytokines including GM-CSF and IL-22 (Coquet et al., 2008; Doisne et al., 2011; Paget et al., 2012). The importance of GM-CSF synthesis by iNKT cells in controlling bacterial infections was identified following *Mycobacterium tuberculosis* infection, although production was not attributed to a particular iNKT cell subset (Rothchild et al., 2014). The requirement for GM-CSF in the response to *S. pneumoniae* was confirmed here, in agreement with previous data indicating the importance of GM-CSF (Brown et al., 2017; Steinwede et al., 2011). Among T cells, early GM-CSF production was specific to NKT17 cells. Although there are other potential sources of GM-CSF (Yamamoto et al., 2014), antibody blocking of GM-CSF was effective in reducing lung *S. pneumoniae* CFU only in wild type mice, but not in *Traj18^-/-^* mice, supporting a role for GM-CSF from NKT17 cells.

We determined that cDC1 cells were one important APC-type for activating innate-like T cells and optimal protection. Mice lacking this subset due to deletion of *Batf3* had a reduced number of lung iNKT and γδ T cells after infection and a decreased percentage of these cells producing IL-17A, with a concomitant reduction in neutrophil recruitment and increased lung CFU. In a model in which *α*GalCer was administered prior to *S. pneumoniae* infection to elicit protective responses, lung cDC1 cells were implicated (Ivanov et al., 2012). Additionally, deletion of the gene encoding CD1d in cDC1 cells did not affect iNKT cell recruitment, but led to a reduction in iNKT cell activation to produce IL-17A and an increased bacterial load. By contrast, γδ T cell cytokine production was not decreased, indicating these cells are dependent on the cDC1 cell type, and perhaps cytokines they produce, but are not dependent on their CD1d-mediated antigen presentation. Other myeloid populations also express CD1d and/or produce activating cytokines such as IL-1β, and therefore one or more of these populations could be responsible for the remaining iNKT cell and γδ T cell activation. This could occur either through the production of cytokines or other proteins, or in the case of iNKT cells, because of CD1d antigen presentation.

In summary, this study identified previously unknown mechanisms by which innate-like T cells, in part due to interactions with cDC1, protect mice from lung infection with *S. pneumoniae*. For iNKT cells, TCR-mediated GM-CSF production by NKT17 cells, which are pre-located largely outside the vasculature, was important. After infection, γδ T cells rapidly left the vasculature and produced cytokines including IL-17A, and while their activation depended on cCD1 cells, it did not depend on their expression of CD1d. Therefore, the signals inducing γδ TCR activation remain unknown. Importantly, all of these T cell and DC populations are relatively infrequent in the lung, emphasizing the importance of rare and specialized cell types in protection from *S. pneumoniae* and potentially other infections. MAIT cells are more abundant in humans than in mice, and MAIT cell numbers increased in nasal biopsies from human subjects challenged with *S. pneumoniae* (Jochems et al., 2019) and IL-17A producing MAIT cells were increased in children with community acquired pneumonia (Lu et al., 2020). Therefore, it is possible that similar mechanisms dependent on innate-like T cells apply to human lung bacterial infections, although with the more prevalent MAIT cells providing immune support.

## Acknowledgements

Supported by the American Lung Association Senior Research Training Fellowship RT-412662 to C.M.C; US National Institutes of Health R01 AI71922, AI105215, AI137230 to M.K.; T32 AI125179 to M.P.M.; R35 HL135756 and R01 AI115053 to J.P.M.; the German Research Foundation SFB1292, CL 419/2-2, CL 419/4-1, and the Research Center for Immunotherapy (FZI) Mainz to B.E.C.; Shared Instrumentation Grant (SIG) Program S10 OD018499 to the Flow Cytometry Core Facility at the La Jolla Institute for Immunology; S10 RR027366 for a FACSAria II cell sorter to Dr. Michael Croft; Tullie and Rickey Families SPARK Awards for Innovations in Immunology to C.M.C.

## Author Contributions

Conceptualization, M.P.M., C.M.C., Z.M., M.K.; Methodology, C.M.C., P.M., Z.M.; Reagents, B.E.C.; Investigation, M.P.M., C.M.C., P.M., N.H., S.C., A.K., S.Z., M.Z., F.T.C., J.P.M.; Writing – Original Draft, M.P.M, C.M.C., M.K.; Writing – Review & Editing, M.P.M. and M.K., Funding Acquisition, M.K.; Supervision, J.P.M, B.E.C. and M.K.

## Materials and Methods

### Animals

Mice were bred and housed under specific pathogen-free conditions in the vivarium of the La Jolla Institute for Immunology (La Jolla, CA). Age and gender matched, male and female mice ages 8-16 weeks were used in all experiments. Inbred C57BL/6J, *Batf3^-/-^,* GM-CSF^-/-^ (*Csf2^-/-^*) and *Itgax*-cre (*Cd11c*-cre) mouse strains were purchased from Jackson Laboratory (Bar Harbor, ME). *Cd1d^f/f^* mice (Birkholz et al., 2015), *Traj18^-/-^* mice (Chandra et al., 2015), and *Langerin*-Cre knockin mice (Zahner et al., 2011) were generated as described previously. Nur77^GFP^ mice were a gift from Kristin A. Hogquist (University of Minnesota, Minneapolis, Minnesota, USA) (Moran et al., 2011). All procedures were carried out under the Association for Assessment and Accreditation of Laboratory Animal Care (AALAC) and approved by the La Jolla Institute for Immunology Institutional Animal Care and Use Committee (IACUC).

### Streptococcus pneumoniae strains

*S. pneumoniae* serotype 3 strain URF918 is a clinical isolate originally from Japan (Kawakami et al., 2003). GFP-*Streptococcus pneumoniae* was generated in URF918 as previously described (Kjos et al., 2015). *S. pneumoniae* serotype 19A isolates, used for in vitro iNKT cell activation experiments, originated from the sites listed in Supplementary Table 1.

### Streptococcus pneumoniae infection

For mouse infections, URF918 was cultured in Todd-Hewitt broth (BD Biosciences) plus yeast extract at 37°C in an incubator at 5% CO_2_, collected at a mid-log phase and washed twice in PBS. For pulmonary infection, mice were anesthetized with isoflurane and elevated on a board. Mice were inoculated with *S. pneumoniae* (1-3×10^6^ colony-forming units in a volume of 50 μl per mouse) by insertion of a pipet tip into the trachea. For calculation of total lung bacterial burden, tissues were collected at day 2 after infection and were homogenized in PBS to assess bacterial burden. Homogenates were inoculated at different dilutions in a volume of 50 μl on 5% sheep blood agar plates (Hardy Diagnostics, Santa Maria, CA) and cultured for 18 h, followed by counting of colonies. Note that the cfu in WT mice varied in different experiments and comparisons are therefore generally only valid within an experiment. For blocking experiments, mice were treated with 10 ug purified NA/LE Rat IgG2a isotype control antibody (BD Biosciences, San Diego, CA) or Ultra-LEAF purified anti-mouse GM-CSF antibody (BioLegend, San Diego, CA) 6 hours prior to infection, as previously described (Brown et al., 2017). Lungs were harvested 16 hours post-infection. For bacterial uptake, lungs were harvested 2 hpi and processed as described for flow cytometry or immunofluorescence.

### Processing of mouse lung tissue for flow cytometry

Mouse lungs were removed and rinsed with RPMI+10% FBS. The lungs were placed in a GentleMacs C tube (Miltenyi Biotec, Bergisch Gladbach, Germany) with 2mL STEMCELL Spleen Dissociation Medium (STEMCELL Technologies, Vancouver, BC, Canada), and homogenized using the Miltenyi GentleMacs dissociator. The cell suspensions were filtered with a 70 μm filter and washed with RPMI+10% FBS. For intracellular cytokine experiments, the cell suspensions were cultured with Golgi-Stop and Golgi-Plug (BD Biosciences) in 2 mL RPMI+10% FBS for 2 hours. Cell suspensions were washed and stained for flow cytometry.

### Discrimination of tissue and vascular resident cells

Mice were anesthetized with isofluorane and injected retro-orbitally with 3 μg of Alexafluor700-labeled anti-CD45 antibody (30-F11), as described previously (Barletta et al., 2012; Reutershan et al., 2005). After 3 minutes, the lungs were removed for processing.

### BrdU incorporation assay

At the time of infection, mice were injected intraperitoneally with BrdU. Lung tissues were collected at 24 hours and stained for surface markers and BrdU according to the manufacturer’s protocol (BD Biosciences).

### Flow cytometry

For staining of cell surface molecules, cells were suspended in staining buffer (PBS, 2% BSA) and stained with fluorochrome-conjugated antibodies at a concentration of 1:200 for 20 min in a total volume of 50 μl. FcγR-blocking Ab anti-CD16/32 (2.4G2) was added to prevent nonspecific binding. Cells were fixed with 2% formaldehyde for 30 minutes on ice. For intracellular cytokine staining, cells were permeabilized with diluted ThermoFisher 10x permeabilization buffer for 5 minutes and stained overnight at 4° C in 1x permeabilization buffer. Cells were washed thoroughly, analyzed in an LSR II flow cytometer (BD Biosciences), and data were processed with Flow Jo software (Tree Star, Ashland, OR).

iNKT cells were stained using tetramers of CD1d loaded with αGalCer (BV421, in house preparation), live/dead yellow (ThermoFisher Scientific), anti-TCRβ-APC-eF780 (H57-597, ThermoFisher Scientific), anti-CD8*α*-BV605 (53-6.7, BioLegend) and anti-CD19-BV605 (1D3, BD Biosciences), anti-CD4-AF700 (GK1.5, BioLegend), anti-ICOS-Pe Cy7 (C398.4A, BioLegend), anti-CD49a-BV711 (HA31/8, BD Biosciences). For intracellular cytokine staining the following antibodies were used: anti-GM-CSF-PE (MP1-22E9, BioLegend); anti-IL-17A (TC11-18H10.1, BioLegend); anti-IFN-γ (XMG1.2, BioLegend). iNKT cell subsets were gated as follows (Supplementary Figure 2): live lymphocytes, singlets, CD8^-^CD19^-^, CD45^+^, Tetramer^+^TCRβ^+^ iNKT cells and separated into NKT1, NKT2, and NKT17 cell subsets based on the following expression profiles: NKT1: CD49a^+^ICOS^-^; NKT2: CD49^-^ICOS^+^CD4^+^; NKT17: CD49a^-^ICOS^+^CD4^-^.

Staining of antigen presenting cells used the following reagents, gating strategy shown in Supplementary Figure 3A: live/dead yellow (ThermoFisher Scientific), anti-CD45-BV786 (30.F11, BD Biosciences); anti-siglecF-BV421 (E50-2440, BD Biosciences); anti-Ly6G-PE (1A8, BioLegend); anti-CD11b-PerCP-Cy5.5 (M1/70, BD Biosciences); anti-CD11c-APC (N418, eBioscience); anti-CD103-BV711 (M290, BD Biosciences); anti-CD24-FITC (M1/69, BD Biosciences); anti-CD64-PE Cy7 (X54-5/7.1, BioLegend); anti-CD86 (GL-1, BD Biosciences). For intracellular cytokine staining, anti-IL-23p19-AF488 (fc23cpg, Invitrogen); anti-IL-1β-APC (NJTEN3, eBioscience). Following selection for single, live, CD45^+^ cells, antigen presenting cells were gated as follows (Supplementary Figure 4A): cDC1: Ly6G^-^SiglecF^-^CD11c^+^CD103^+^; cDC2: Ly6G^-^SiglecF^-^CD11c^+^CD103^-^ CD11b^+^CD64^-^CD24^+^; moDC: Ly6G^-^SiglecF^-^CD11c^+^CD103^-^CD11b^+^CD64^+^CD24^+^; alveolar macrophages: SiglecF^+^Ly6G^-^CD11c^+^; neutrophils: Ly6G^+^CD11b^+^.

MAIT cells were stained using 5-OP-RU-MR1-Tetramer or 6-FP-MR1-Tetramer (NIH Tetramer Core). Single cell suspensions were stained with tetramers for 45min at room temperature. Cells were washed twice with staining buffer then incubated with antibodies for further surface staining. γδ T cells and MAIT cells were gated as follows: live lymphocytes, singlets, CD11b^-^CD19^-^CD45^+^γδTCR^+^ (anti-TCR γδ-FITC, clone GL3; γδ T cells); live lymphocytes, singlets, CD11b^-^CD19^-^CD45^+^γδ TCR^-^TCRβ^+^5-OPRU Tetramer^+^ (MAIT cells). Cytokine staining/antibodies described above.

### Confocal microscopy

For bacterial uptake, Langerin-YFP^+^ mice were infected with 10^7^ GFP- *S. pneumoniae* as described above and euthanized 2 hpi. Lungs were inflated with low-melting point agarose and processed into 300 µm sections with a Vibratome. Sections were individually placed in a multi-well plate, washed, and fixed with paraformaldehyde, washed, and embedded in Prolong Glass anitfade (ThermoFisher Scientific) under #1.5 coverslip.

Nur77^GFP^ mice were inoculated with *S. pneumoniae* as described above and euthanized after 14h. Lungs were inflated with low-melting point agarose and processed into 300 µm sections with a Vibratome. Sections were individually placed in a multi-well plate, blocked with FcγR-blocking Ab (2.4G2) in PBS for 1 hour at RT, and then stained overnight at 4° C on a rocker in a 500 µl volume. PE-labeled CD1d tetramer loaded with αGalCer was used at 1:200 dilution, and anti-CD31-BV421 antibody (clone 390, BioLegend) was added at 1:100 dilution. After washing, sections were fixed with paraformaldehyde, washed and embedded in Prolong Glass anitfade (ThermoFisher Scientific) under #1.5 coverslip.

Slides were imaged with ZEISS LSM780 confocal microscope using 40x/1.3NA EC Plan-Neofluar oil objective. Fluorescence of BV421, GFP, and PE was excited with 405 nm, 488 nm, and 561 nm laser lines, and emitted signals were collected on spectrally tuned bialkali PMT and GaAsP detectors. Single-stained controls were used to ensure no bleed-through between channels. Unloaded tetramers were used as a specificity control. Pixel size was set to 115 nm and Z-stacks were acquired with a 3 um step size using tile scan function. Average field of view was 370×370×70 µm, and 3 regions were acquired per sample. Images were loaded into Imaris (9.7.2, Oxford Instruments) and 3D views were used to examine the location of cells positive for GFP and tetramer staining in respect to CD31-labeled vasculature. Channel brightness was adjusted to improve contrast in the same manner across all images. Orthogonal top views were created using Snapshot function in Imaris and figures were arranged in Photoshop.

### *in vitro* iNKT cell activation and ELISA

Preparation of bacterial lysates and cell-free antigen-presentation assays have been described previously (ref). Briefly, bacterial sonicates were incubated for 24 hours in microwells coated with soluble mouse CD1d or purified human PBMCs. After washing, 1 x 10^5^ mouse 1.2 V*α*14i NKT cell hybridoma cells were added to the wells for 20 to 24 h. Mouse IL-2 was measured in the supernatants by enzyme-linked immunosorbent assay (BioLegend, BD Biosciences).

### Statistical Analysis

Graphs and statistical analyses were generated with Prism 7 and 9 software (Graphpad, 2016).

