## Supplementary Figures for "Specialized subsets of innate-like T cells and dendritic cells protect from lethal pneumococcal infection in the lung"

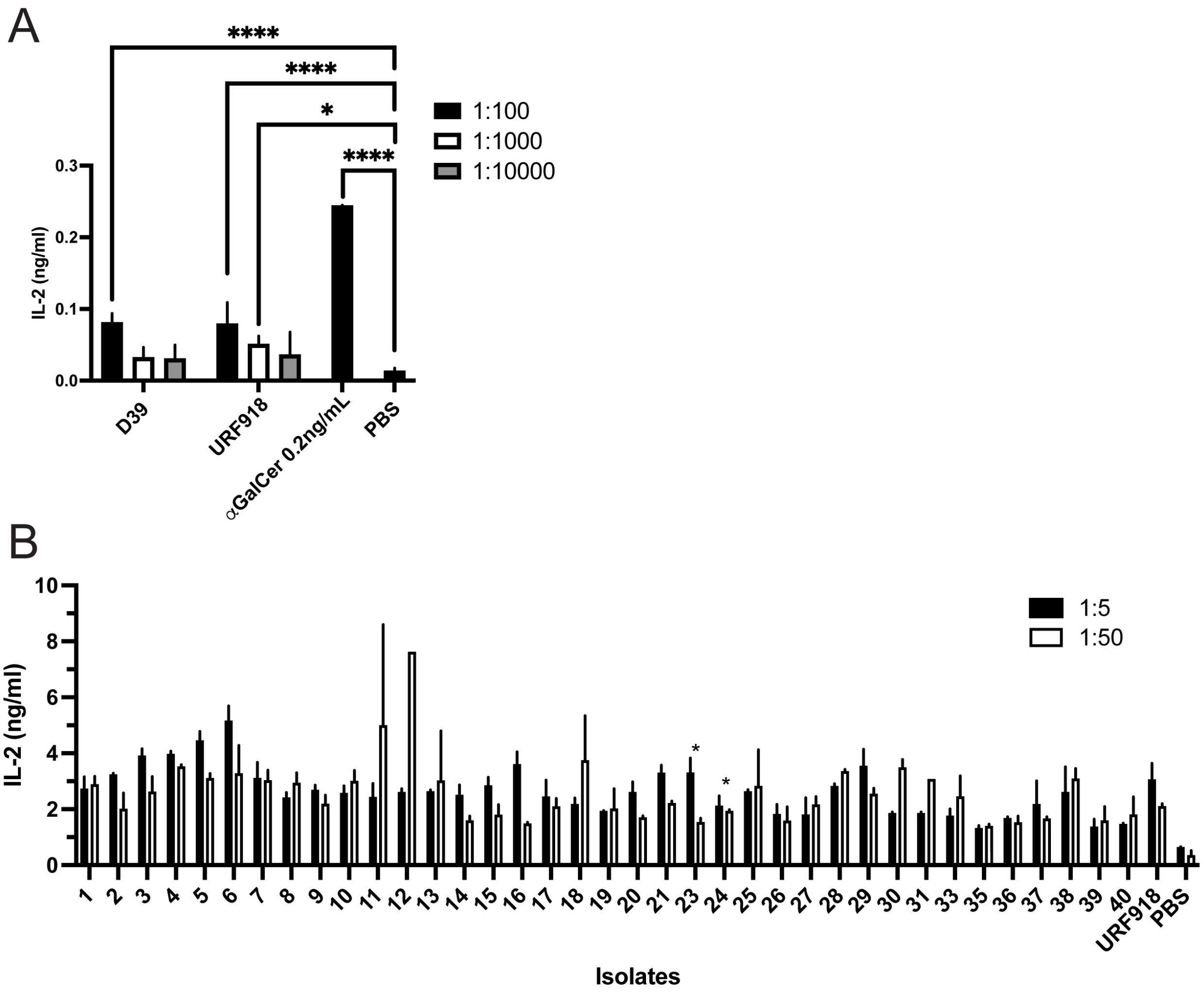


**Supplementary Figure 1: Clinical Isolates of *S. pneumoniae* activate iNKT cells *in vitro*.** IL-2 production by ELISA following CD1d-coated plate assay testing *S. pneumoniae* clinical isolates. A) D39 and URF918 sonicates with α-GalCer 0.2 ng/mL. Statistical significance assessed via 2-way ANOVA with Dunnett's multiple comparisons test, **** = P < 0.0001; * P = 0.0173 (Dunnett's). B) Sonicates of serotype 19A clinical isolates with specified dilutions. Invasive strains denoted by *. Representative of two independent experiments; error bars represent technical replicates (2). Statistical significance assessed via 2-way ANOVA with Dunnett's multiple comparisons test, comparing each sample to PBS. P values listed in supplementary table 1. URF918 vs PBS: * P = 0.0319 (Dunnett's).

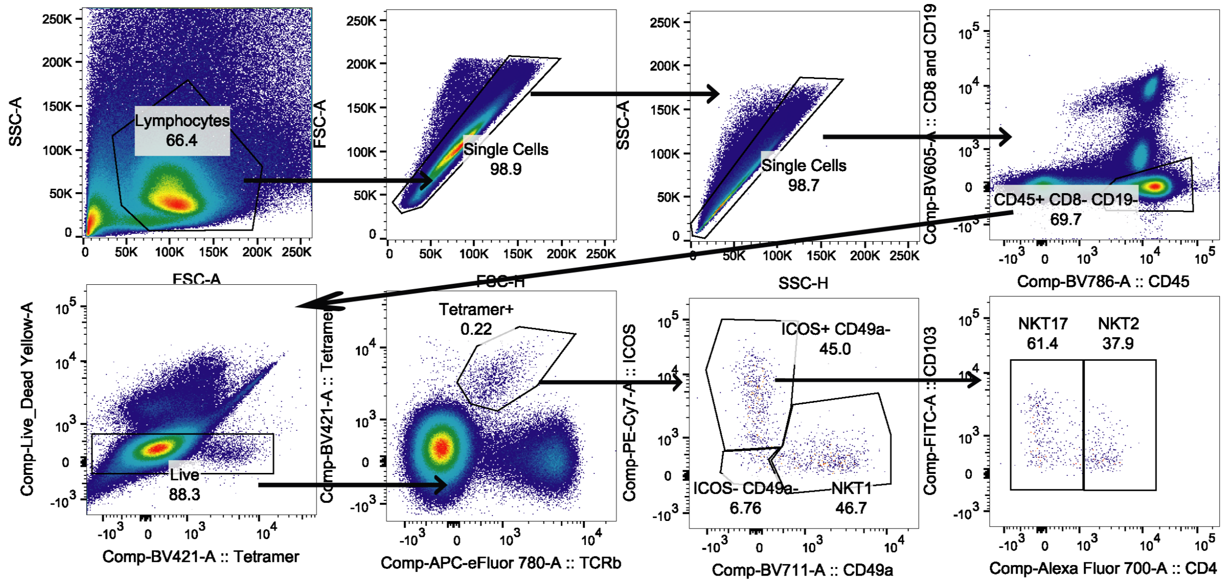


**Supplementary Figure 2: Gating Strategy.** Gating strategy for iNKT cell subsets in uninfected lung.

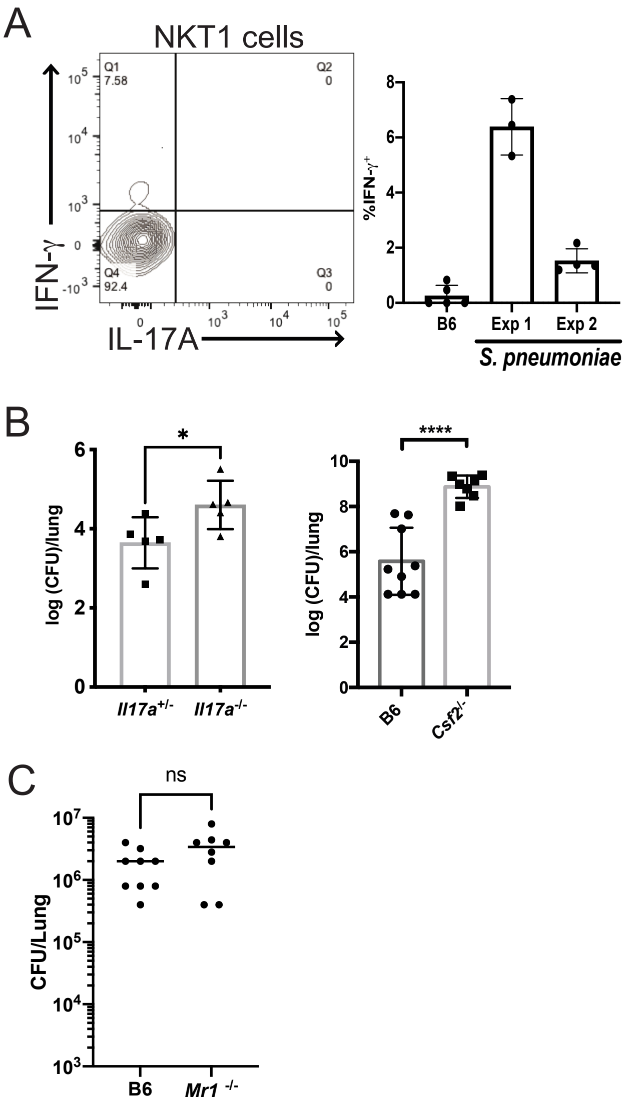


**Supplementary Figure 3: Innate-like T cell responses to *S. pneumoniae*.** A) Representative flow cytometry plot showing IFN-γ production by NKT1 cells (left) and quantitation from two different experiments on right. N = 3-5 mice per group. B) *Il17a*- and *Csf2*- deficient mice were infected with URF918 and bacterial burden within the lung was assessed at 48 hours (left) and 18 hours (right) post-infection. IL-17A-deficient mice: 5 mice per group, representative of one of 2 independent experiments, unpaired t test. *Csf2*$^{-/-}$ mice: 7-9 mice per group, data compiled from 2 independent experiments, unpaired t test. C) Bacterial burden in the lung of C57BL/6J (B6) mice and *Mr1*$^{-/-}$ mice 2 dpi with URF918.

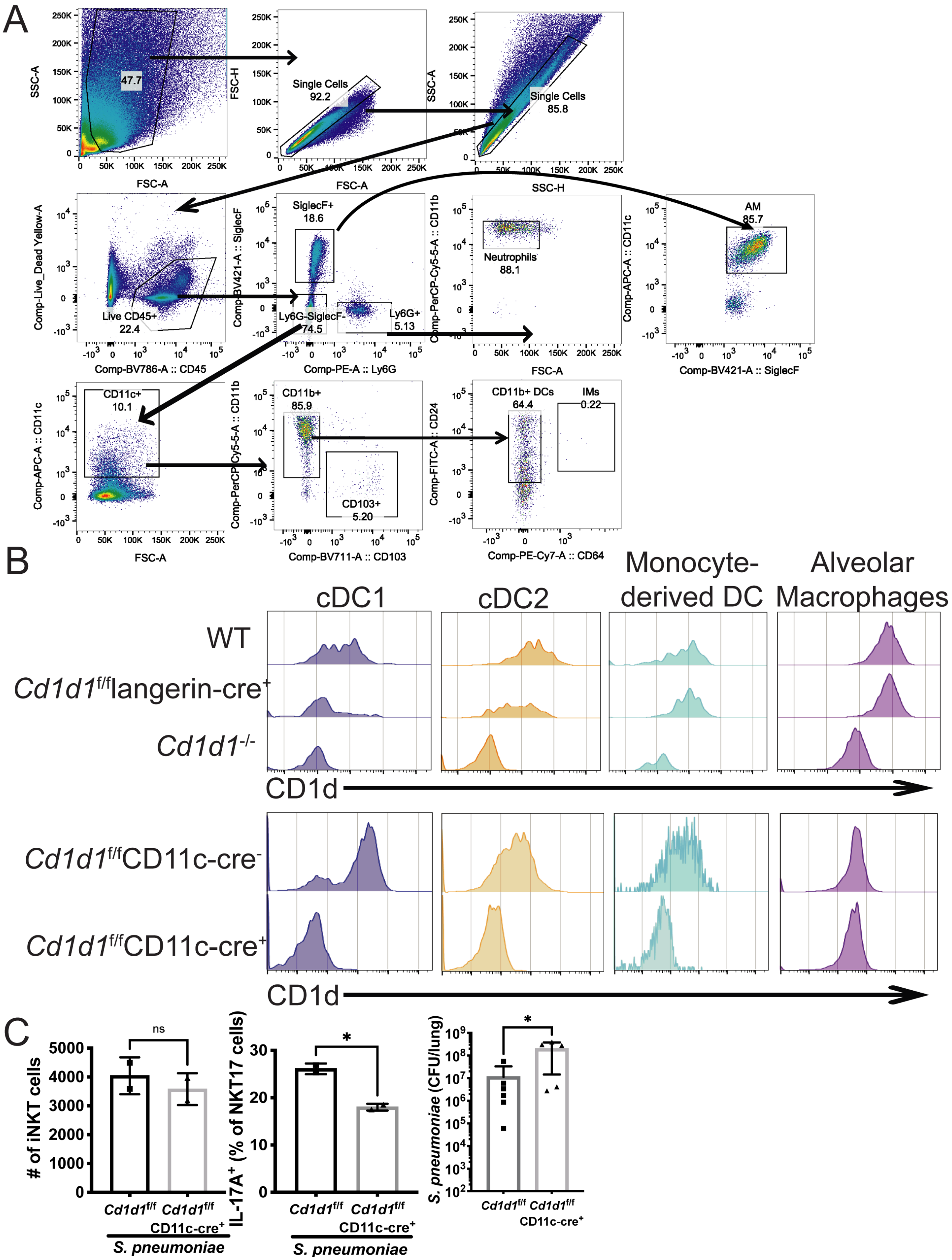


**Supplemental Figure 4: CD1d expression by lung antigen presenting cells.** A) Gating strategy of antigen presenting cells and neutrophils in uninfected lung. B) CD1d expression assessed by flow cytometry in lung APCs of C57BL/6J, *Cd1d1*$^{f/f}$ x *Langerin*-Cre$^{+}$ mice, *Cd1d1*$^{-/-}$ mice, and *Cd1d1*$^{f/f}$ x *CD11c*-Cre$^{+}$ mice. Alveolar macrophages express CD11c but expression of CD1d was not affected by the CD11c-cre. C) Analysis of *Cd1d1*$^{f/f}$-CD11c-cre$^{+}$ mice following infection with URF918. Left: Total iNKT cell numbers at 15 hours post-infection. Middle: IL-17A production by NKT17 cells at 15 hours after infection. Right: Bacteria loads 2 days after infection (unpaired t test).

**Supplementary Table I. Serotype 19A sites of origin.**

| Isolate ID | MLST | Site of Origin | Serotype | Plate no./ Isolate no. | Dunnett's multiple comparisons test | |
|---|---|---|---|---|---|---|
| | | | | | Summary | Adjusted P Value |
| 06AR0388 | 695 | Blood | 19A | 1 | * | 0.0109 |
| 06AR0474 | 1925 | Blood | 19A | 2 | * | 0.0261 |
| BMC0124C | 199 | Nasopharynx | 19A | 3 | *** | 0.0009 |
| BMC0039P | | Nasopharynx | 19A | 4 | **** | <0.0001 |
| BMC0008C | | Nasopharynx | 19A | 5 | **** | <0.0001 |
| BMC0196C | | Nasopharynx | 19A | 6 | **** | <0.0001 |
| 06AR0478 | 695 | Blood | 19A | 7 | ** | 0.0027 |
| BMC0014C | | Nasopharynx | 19A | 8 | * | 0.0207 |
| 08AR0009 | | Pleural fluid | 19A | 9 | ns | 0.0603 |
| 05AR0302 | 415 | Pleural fluid | 19A | 10 | * | 0.012 |
| 08AR0026 | 695 | pleural fluid | 19A | 11 | **** | <0.0001 |
| BMC0141C | | Nasopharynx | 19A | 12 | **** | <0.0001 |
| BMC0213C | | Nasopharynx | 19A | 13 | ** | 0.0098 |
| 07AR0013 | 667 | Blood | 19A | 14 | ns | 0.2422 |
| BMC0401C | 3637 | Nasopharynx | 19A | 15 | ns | 0.0953 |
| BMC0236C | 2083 | Nasopharynx | 19A | 16 | * | 0.0378 |
| 06AR0400C | 695 | Blood | 19A | 17 | ns | 0.1148 |
| BMC0359C | | Nasopharynx | 19A | 18 | ** | 0.0049 |
| BMC0200C | | Nasopharynx | 19A | 19 | ns | 0.3099 |
| 07AR0061 | 199 | Blood | 19A | 20 | ns | 0.1734 |
| 03AR0753 | 199 | Pleural fluid | 19A | 21 | * | 0.0141 |
| 10AR0013 | 320 | Invasive | 19A | 23 | ns | 0.0634 |
| 05AR156 | 320 | Invasive | 19A | 24 | ns | 0.2659 |
| 08AR0090 | 199 | Unknown | 19A | 25 | * | 0.0158 |
| 09AR0097 | 199 | Unknown | 19A | 26 | ns | 0.6152 |
| 06AR0008 | 199 | Unknown | 19A | 27 | ns | 0.3003 |
| 04AR0776 | 199 | Unknown | 19A | 28 | ** | 0.0025 |
| 09AR0029 | 695 | Unknown | 19A | 29 | ** | 0.0032 |
| 07AR0143 | 695 | Unknown | 19A | 30 | * | 0.0211 |
| 10AR0019 | 695 | Unknown | 19A | 31 | ns | 0.0536 |
| BMC0196C | 695 | Unknown | 19A | 33 | ns | 0.2073 |
| 06AR0239 | 667 | Unknown | 19A | 35 | ns | 0.9568 |
| 08AR0042 | 667 | Unknown | 19A | 36 | ns | 0.7419 |
| 02AR0813 | 667 | Unknown | 19A | 37 | ns | 0.3648 |
| 05AR0276 | 320 | Unknown | 19A | 38 | ** | 0.0089 |
| 06AR0243 | 320 | Unknown | 19A | 39 | ns | 0.8678 |
| 09AR0039 | 320 | Unknown | 19A | 40 | ns | 0.6928 |
